# Exploration of DPP-IV inhibitory peptide design rules assisted by deep learning pipeline that identifies restriction enzyme cutting site

**DOI:** 10.1101/2022.06.13.495896

**Authors:** Changge Guan, Jiawei Luo, Shucheng Li, Zheng Lin Tan, Yi Wang, Haihong Chen, Naoyuki Yamamoto, Chong Zhang, Yuan Lu, Junjie Chen, Xin-Hui Xing

## Abstract

Mining of anti-diabetic dipeptidyl peptidase IV (DPP-IV) inhibitory peptides (DPP-IV-IPs) is currently a costly and laborious process. Due to the absence of rational peptide design rules, it relies on cumbersome screening of unknown enzyme hydrolysates. Here, we present an enhanced deep learning (DL) model called BERT-DPPIV, specifically designed to classify DPP-IV-IPs and exploring their design rules to discover potent candidates. The end-to-end model utilizes a fine-tuned bidirectional encoder representations (BERT) architecture to extract structural/functional information from input peptides and accurately identify DPP-IV-Ips from input peptides. Experimental results in benchmark dataset showed BERT-DPPIV yielded state-of-the-art accuracy of 0.894, surpassing the 0.797 obtained by sequence-feature model. Furthermore, we leverage the attention mechanism to uncover that our model could recognize restriction enzyme cutting site and specific residues that contribute to the inhibition of DPP-IV. Moreover, guided by BERT-DPPIV, proposed design rules of DPP-IV inhibitory tripeptides and pentapeptides were validated and they can be used to screen potent DPP-IV-IPs.

## Introduction

Due to the side effects and the requirement of injection of commercial dipeptidyl peptidase IV (DPP-IV) inhibitors, which are used to treat approximately 537 million patients with type II diabetes, it is crucial to develop new DPP-IV inhibitory drugs and functional foods^1,2,3^. Peptides with DPP-IV inhibitory activities are a promising class of oral hypoglycemic without adverse effects, which are derived from products of enzymatically hydrolyzed edible animal, plant, and macroalgal proteins^1^.

Current approaches for the discovery of DPP-IV inhibitory peptides (DPP-IV-IPs) are known to be labor- and cost-intensive. For instance, conventional enzymatic hydrolysate screening method requires extensive separation and purification, mass spectrometric identification and revalidation of peptide synthesis^2,3^. In addition, the lack of design rules and effective characterization methods make it challenging to design DPP-IV-IPs and synthesis peptide libraries for DPP-IV-IPs screening^4^. These problems have limited the discovery of efficient DPP-IV-IPs. Therefore, it is an important task to develop a high-throughput method capable of rapidly identifying effective DPP-IV-IPs that can be used in functional foods or medicine innovation^1,2,5^.

Recently, powered by artificial intelligence, large language models (LLMs) based on natural language processing (NLP) have been successful used to solve biological problems. Therefore, computation tools complement experimental studies in DPP-IV-IPs discovery process. Over the past decade, several computational approaches have been used for the discovery of DPP-IV-IPs including quantitative structure-activity relationship method^6–9^ and machine learning (ML)^7,10–13^. The ML methods mainly include support vector machine and random forest^14–16^. However, such methods reported to date have some major limitations leading to poor performance: (1) the performance of these models is heavily dependent on the quality of the features extracted by feature engineering, (2) they are poorly able to represent amino acid sequences, and (3) they fail to capture information hidden in the amino acid sequence itself, which renders these models inefficient. Furthermore, classifiers for DPP-IV-IPs developed to date have not been verified experimentally due to low accuracy of model which will result in high cost of experimental verification.

Deep neural networks with advanced architecture can overcome these limitations, and have a capability of automating feature learning to extract discriminating feature representations with minimal human effort^17^. In particular, DL methods, e.g., long-short term memory, which is based on NLP and regarded amino acid sequence as natural language, is a promising method in identifying functional peptides for extraction of discriminative feature representation, such as anti-microbial peptides^18–21^. However, DL-based models have been criticized for their interpretability and unexplainable characteristics, which are referred to as a ‘black box’. Moreover, limited volume of data is a great challenge to training DL models from scratch.

In this article, we present a novel peptide language model (PLM)-based DL model, named BERT-DPPIV, to offer a promising solution to address the challenges of DPP-IV-IPs screening and design. BERT-DPPIV combines a pre-training process based on unlabeled protein data and a NLP BERT model with an attention mechanism to identify DPP-IV-IPs. By learning the discrimination task, BERT-DPPIV was trained to automatically learn the characteristics and features contained in the original peptide sequences and to distinguish and represent amino acid sequences in high-dimensional spaces, and achieved state-of-the-art.

Benefiting from the attention mechanism, BERT-DPPIV has good interpretability, which is more advantageous than existing ML algorithms for DPP-IV-IPs identification. According to the attention analyses, we have found that our model can learn tertiary structure property of peptide except for physicochemical properties. Particularly, we have found that BERT-DPPIV can identify cleavage sites of DPP-IV enzymes on the substrate polypeptides, which was first reported, and suggested that NLP DL model may be used to analysis enzyme cleavage site.

The accuracy of BERT-DPPIV was validated by wet laboratory experiment based on measurement of half maximal inhibitory concentration (IC_50_) of synthetic peptides predicted to be DPP-IV-IPs by BERT-DPPIV, and the result suggest that the prediction accuracy is in align with experimental data, thus extending the computational model to practical applications. By combining both the PLM model and biological assays, we have proposed a novel DPP-IV inhibitory pentapeptide design strategy based on the dipeptide repeat unit X-proline to provide a new idea for the design of DPP-IV-IPs. This design strategy has addressed the bottleneck in DPP-IV-IP discovery and demonstrated that *in silico* evaluation method can aid peptide drug development.

## Results

### Overview of BERT-DPPIV

BERT-DPPIV is a PLM-based DL framework designed for mine novel DPP-IV-IPs based on their amino acid sequence (Fig 1). To realize this, the framework consists of four main components: pre-training, fine tuning, analysis, and model validation and application. During the pre-training procedures, a total of 556,603 protein sequences from databases including UniProt, SWISS-PROT, TrEMBL, and PIR-PSD were used to pre-trained 12 layer of BERT-based language models. We employed three different word segmentation approaches, i.e., kmer = 1, kmer = 2, kmer = 3 to generate three pre-trained BERT models. Two major tasks, i.e., masked language model (MLM) and next sentence prediction (NSP) involved in the pre-training procedures for capturing word-level and sentence-level representations and learning the common features of protein sequences. The fine-tuning process aimed to construct PLM model specifically for the task of identifying DPP-IV-IPs.

**Fig. 1.**
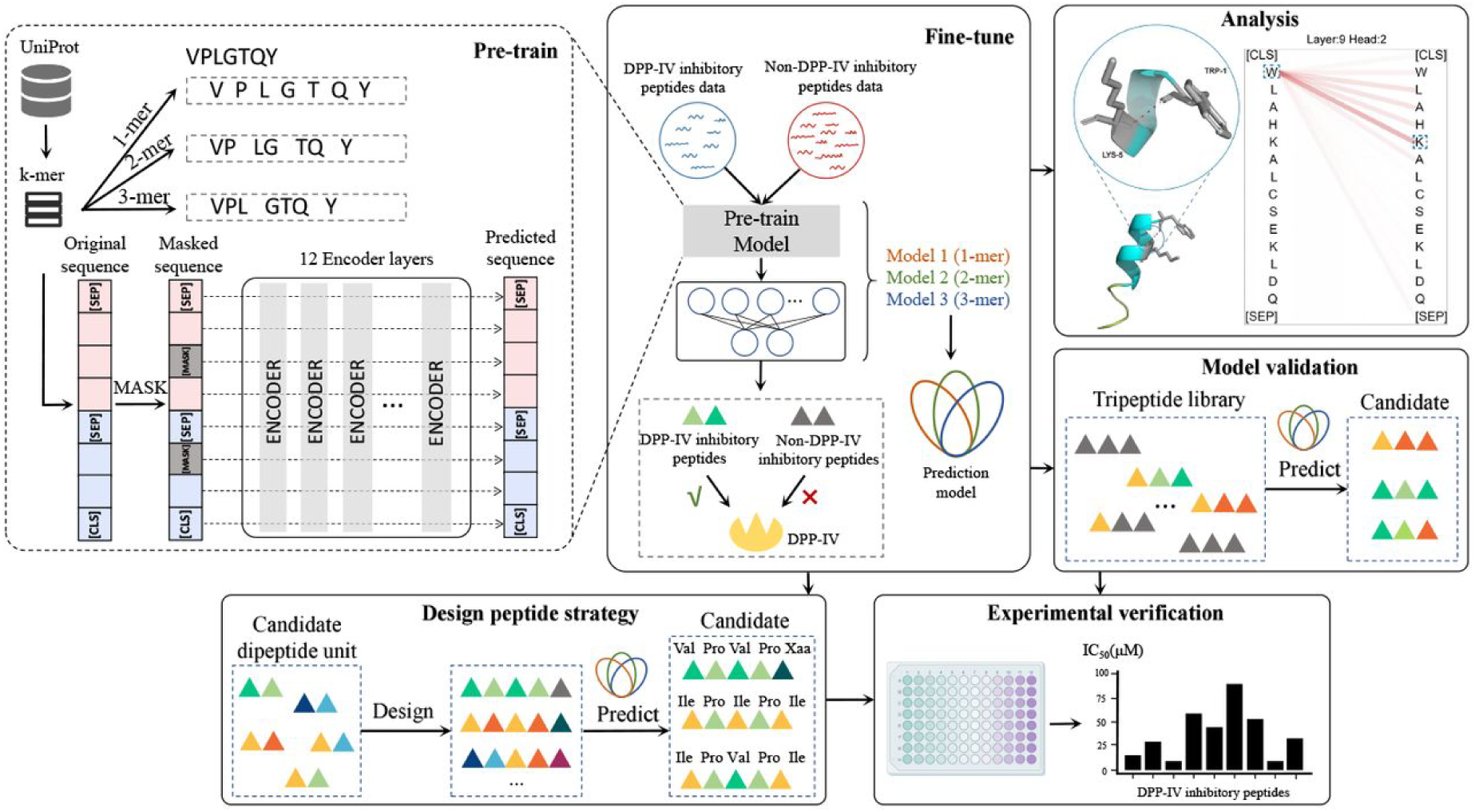
Schematic representation of study workflow. We first collected sequences to pre-train a BERT model (upper left). We then used the DPP-IV-IP dataset for fine-tuning the pre-trained BERT model to obtain three kinds of PLMs. The three models were combined to construct the BERT-DPPIV (upper middle). To determine what the model had learned, we analyzed and visualized the model’s attention (upper right, analysis module). We then used BERT-DPPIV to screen tripeptides that may inhibit DPP-IV to illustrate the performance of our model inhibition (upper right, model validation module). After the model screening, the functional peptides screened out by the model were verified by biological experiments (lower right). Having demonstrated the practical applicability of our model, we proposed a design strategy for DPP-IV-IPs based on dipeptide repeat units and demonstrated the feasibility of this strategy using models and experiments (lower left).

The framework’s analysis component involved evaluating the performance and capabilities of BERT-DPPIV. Various metrics were conducted to assess the model’s ability to predict and identify DPP-IV-IPs accurately. Three models with different kmers in the test set showed the highest accuracy (Acc) of 0.891 was achieved for kmer = 1, followed by 0.887 for kmer = 2, and 0.842 for kmer = 3 and BERT-DPPIV combining three models achieved state-of-the-art, which resulted in an improved Acc of 0.894 (Supplementary Table. 1).

Our models (kmer=1, kmer=2, and kmer=3) outperform the first sequence-based iDPPIV-SCM model, as evidenced by significant improvements in Acc and Matthews correlation coefficient (MCC) respectively (Supplementary Table. 1). These results highlight the ability of our models to extract more valuable information from peptide sequences. To further compared with sequence-based physicochemical property SVM model and the sequence- and structure-based StackDPPIV model. Our model (kmer=1) achieves the same accuracy as the best model StackDPPIV and BERT-DPPIV outperforms it. These results suggest that our models are able to extract physicochemical properties or structure information and BERT-DPPIV can make good use of information from different models. Moreover, the visualization of the model demonstrated these findings further.

Finally, the model was validated and applied to real-world scenarios, providing a tool for mining and discovering novel DPP-IV-IPs based on their amino acid sequences. Overall, we demonstrate the capability of BERT-DPPIV to predict important parameters for DPP-IV-IPs to guide DPP-IV-IPs screening, i.e., (1) inhibitory activity of peptides, (2) physicochemical properties of peptides sequences, (3) structural properties from peptide sequences, and (4) capture cleavage site information. Furthermore, we have also shown the application of BERT-DPPIV in guiding the design of DPP-IV-IPs in addition to peptides screening. Pentapeptides were designed, and their DPP-IV inhibitory activity was predicted and verified.

### Sequence representation and its properties learned by BERT-DPPIV

To investigate the information captured by our models, we selected kmer = 1 that exhibit the highest Acc among our three kmer models for further study. First, we analyzed the representation of the peptide sequences learned by the pretrained BERT, as prior studies have shown that BERT models can effectively learn the representation of the sequence^18–21^. The vector representations of the peptide sequences during the training process of the model were extracted and projected to 2D by t-distributed stochastic neighbor embedding (t-SNE) for visualization (Supplementary Fig. 1), which showed that the model was able to identify DPP-IV-IPs by learning the vector representation of peptides.

In addition, we have illustrated the ability of our model to learn physicochemical properties of peptides by visualizing the distribution of amino acid and peptide representations. The vector representation learned by the model was well able to distinguish amino acids with different properties (Fig. 2a). We then used the modlAMP package^22^ to extract 10 kinds of physicochemical property information from peptide sequences. Based on peptide sequence physicochemical properties, the vector representation of it was then analyzed. The vector representations learned by the model were able to distinguish among sequences with different physical and chemical properties (Fig. 2b,c, Supplementary Fig. 2). To this point, we have shown that the PLM can learn the physicochemical properties of the peptide sequences.

**Fig. 2.**
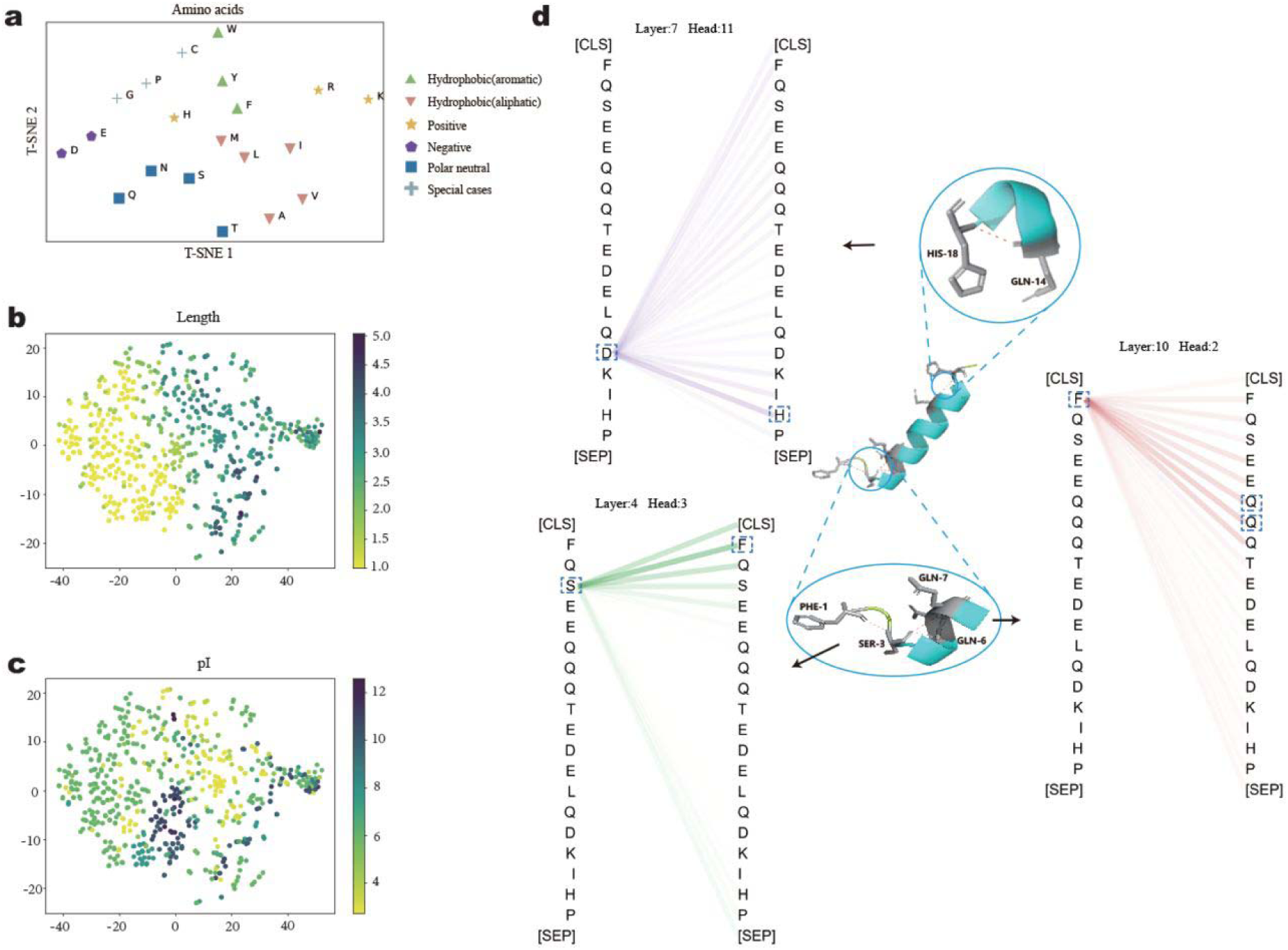
Sequence representation visualization and attention visualization of structural information learned by the model. A vector representation of the peptide sequence and of individual amino acids was extracted from the model and analyzed by t-SNE. The amino acid representation learned by the model (a) and physicochemical properties contained in model-learned representations of peptide sequences (b and c) are shown. Here, we show how the inner workings of the model’s attention heads can be used to analyze a single peptide in more detail. Each attention head performs internally an all-against-all comparison to compute weighted sums for each token relative to all other tokens in the sequence. High scores indicate that the model learned to put more weight on certain residue pairs (upper left, lower left, and lower right). The structure of the peptide shows that Ser-3 can interact with Phe-1, Gln-6, and Gln-7 and that Gln-14 can interact with His-18. When visualizing the attention weights for these sites, we observed that some of the attention heads can capture this interaction information.

### Capturing the structural information and attention patterns generated by the models

The attention mechanism provides a way to reveal ‘black-box’ of DL models and features learned by it^23,24^. In this section, we investigated the capability of our model to learn structural information about peptide sequences with the help of an attention mechanism, which is a technology that mimics cognitive attention. First, the structure of the peptides with a length ≥ 5 amino acids from positive samples was predicted by APPTEST software^25^, and 12 kinds of peptides with α-helix secondary structure were found (Supplementary Fig. 3). Three peptides were randomly selected from among these peptides for further structural confirmation using AlphaFold2^26^. The presence of α-helical structures was confirmed.

Then, the attention of the model for each head at each layer of the model was visualized (12 layers, 12 heads per layer) (Supplementary Fig. 4). As shown in Fig. 1, Fig. 2d, and Supplementary Fig. 5, we observed that the attentions focus on the interacting sites of peptides, suggesting that this model was able to capture the structural information of peptide sequences. Furthermore, analysis of 144 attention mechanisms (Supplementary Fig. 4) showed that several learning mechanisms existed in the model, which include previous-word attention patterns (Supplementary Fig. 6a), next-word attention patterns (Supplementary Fig. 6b), delimiter-focused attention patterns (Supplementary Fig. 6c), specific-word attention patterns (Supplementary Fig. 6d,e), and related-word attention patterns (Supplementary Fig. 6f).

### Attentions of BERT-DPPIV divert to DPP-IV cleavage sites and proline

In most cases, the attention learned by BERT-DPPIV focused on the second and third positions of the N terminus of the polypeptide sequences (Supplementary Fig. 6d-g), which are the cleavage sites of DPP-IV^4^. This result suggested that our model can capture information about DPP-IV cleavage sites by learning the sequence of polypeptide substrate.

To further illustrate the capability of cleavage site prediction, we statistically analyzed the attention sites of the sequences of training set (Fig. 3a, b). These results suggested that the attention of our model focused on the cleavage sites of DPP-IVs. Furthermore, a comparison of the changes in attention positions of the model before and after fine-tuning have supported that our model could capture cleavage site information of DPP-IVs.

**Fig. 3.**
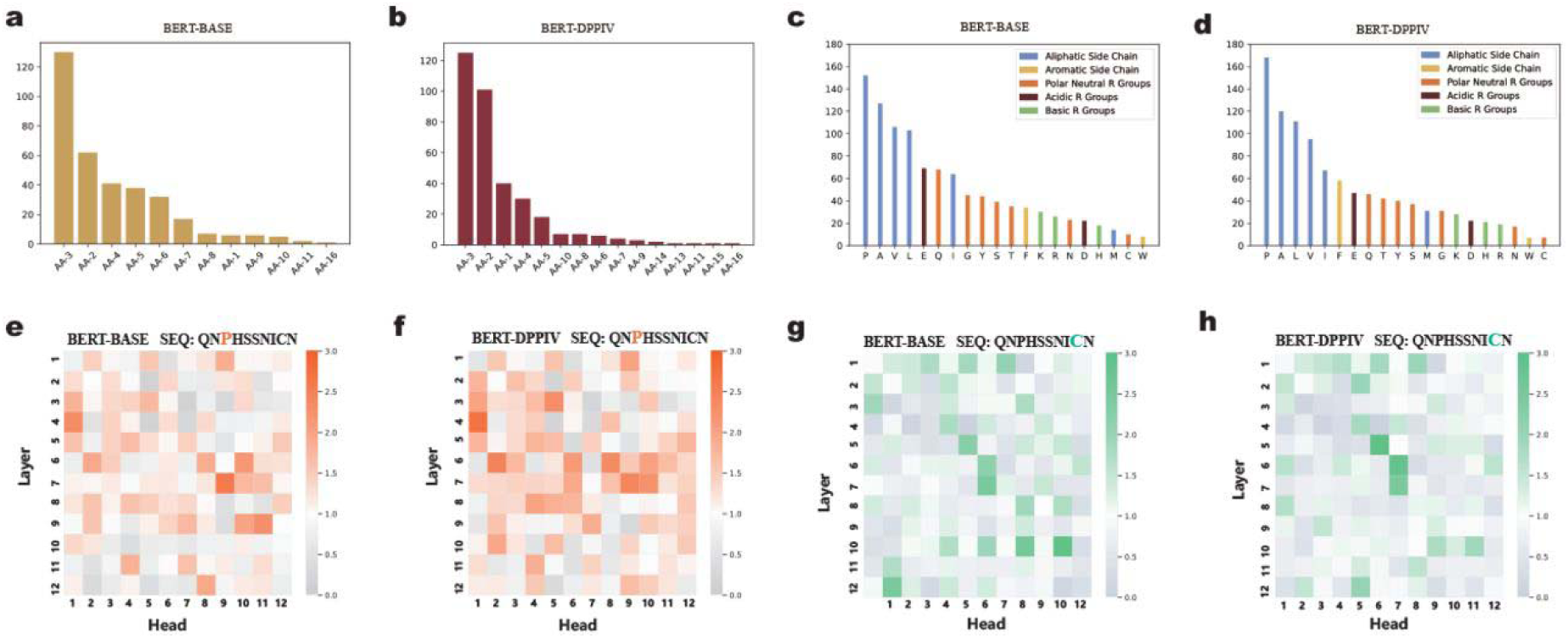
Visualization of the position and importance of amino acid species. To study the importance of position and amino acid species, attention to positions and amino acids was statistically analyzed. BERT-BASE and BERT-DPPIV represent the models before and after fine-tuning, respectively. The statistical importance of the position of peptide sequence according to the attention of the model is shown (a and b). The statistical importance of amino acid species according to the attention of the model is shown (c and d). The attention of the model to proline and cysteine in the peptide sequence was visualized (e, f, g, and h).

We have further investigated the capability of our model to analyze the importance of each amino acid in the sequences. Statistical analyses have suggested that proline is the amino acid of major concern in our model, whereas cysteine is of the least concern. Increase in attention toward proline and decrease in attention toward cysteine were observed (Fig. 3c, d).

To investigate the importance of proline and cysteine in these amino acid sequences, we have visualized the attention of our model for each amino acid in a randomly selected sequence from α-lactalbumin, “QNPHSSNICN”. We observed differences in all amino acids after training, which suggested that the model had directed its attention toward important amino acids, e.g., proline (which is an important amino acid for DPP-IV inhibition), whereas less important amino acids, e.g., cysteine, received less attention (Fig. 3e-h). To illustrate the accuracy of this result, we have statistically analyzed the proportion of our model’s attention that was focused on each of the 20 standard amino acids (Fig.4) and compared it to the proportion of pre-train model’s attention (Supplementary Table 2). These results demonstrated that our model efficiently captured the amino acid types that play important roles in the amino acid sequences of a peptide.

**Fig. 4.**
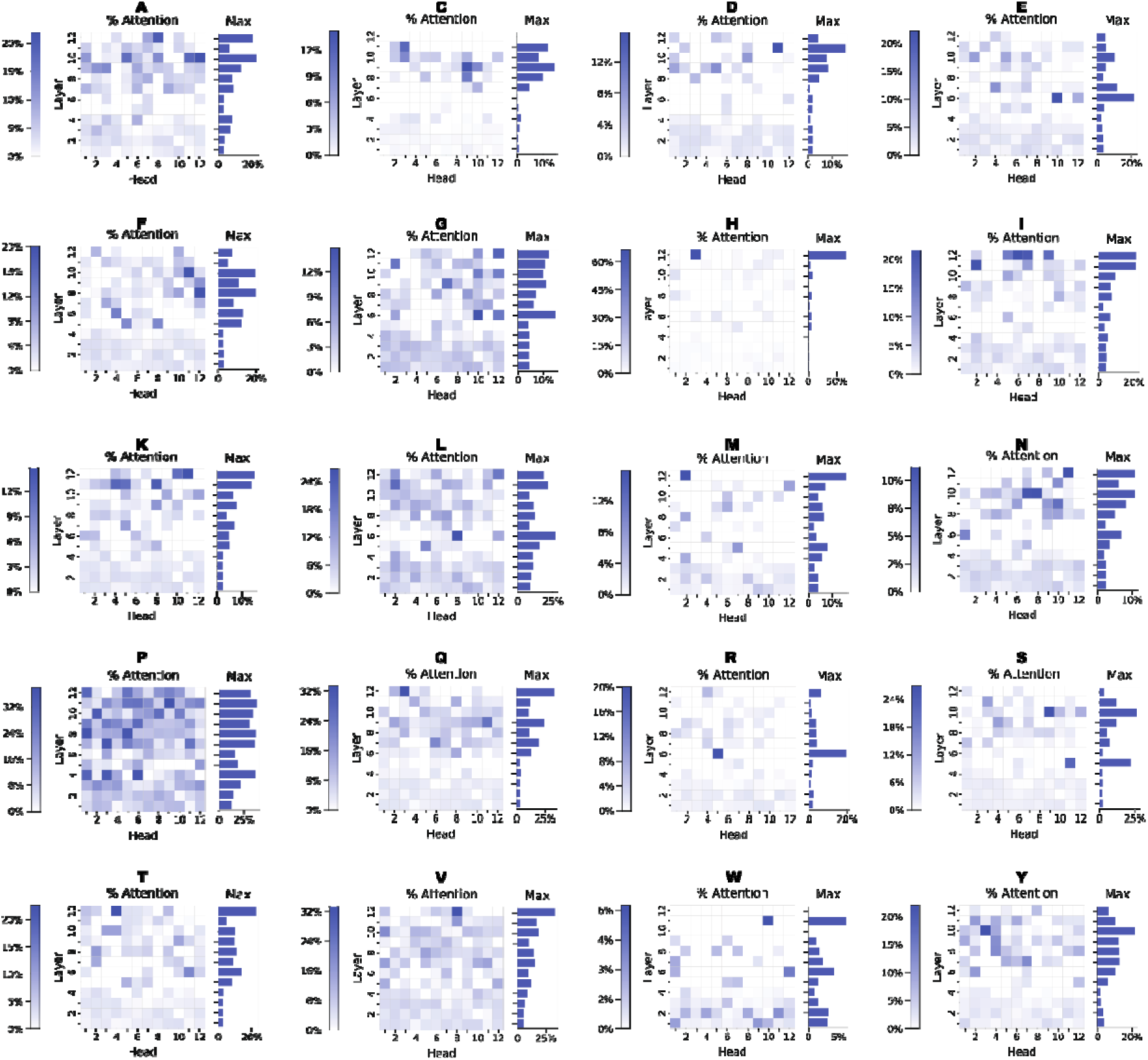
Percentage of each attention head that is focused on the 20 natural amino acids.

### Application of BERT-DPPIV for DPP-IV inhibitory tripeptide screening

To illustrate the usefulness of BERT-DPPIV in mining of DPP-IV-IPs which can be used as anti-diabetic drugs, we applied our model for the screening of DPP-IV inhibitory tripeptides. Prior work conducted by one of the authors (C.G.) classified DPP-IV inhibitory dipeptides into five classes, and, from one class of peptides, represented by VPX and IPX (where X represents one of the 20 amino acids), isolated nine tripeptides with efficient human DPP-IV (hDPP-IV) inhibitory activity^27^. However, it was unknown whether other kinds of tripeptides can efficiently inhibit hDPP-IV. In this section, one class of tripeptides that contains WPX, WAX, WRX, and WVX from five classes previously classified was selected for screening.

We predicted the DPP-IV inhibitory activity of the 80 kinds of tripeptides with our models and found that all these tripeptides are expected to possess DPP-IV inhibitory activity (Supplementary Table 3). Then, these tripeptides were chemically synthesized, and the inhibitory activity was verified experimentally based on the IC_50_, which is inversely correlated to inhibitory activity. IC_50_ were detected from 72 out of the 80 kinds of tripeptides, and the DPP-IV inhibitory activity of each of these tripeptides was confirmed (Table 1). The Acc achieved in this study was 90%, which is consistent with our prediction (Supplementary Table 1). The top three tripeptides for DPP-IV inhibitory activity were WAW, WAY, and WPN with IC_50_ of 103.66 µM, 117.40 µM, and 128.59 µM, respectively. This result demonstrated that our model could predict DPP-IV-IPs with 90% accuracy.

**Table. 1.** Experimental characterization of tripeptides that inhibit DPP-IV

| Peptide | IC<sub>50</sub>, μM | Peptide | IC<sub>50</sub>, μM | Peptide | IC<sub>50</sub>, μM | Peptide | IC<sub>50</sub>, μM |
| --- | --- | --- | --- | --- | --- | --- | --- |
| WRM | 896.67 | WPE | 353.02 | WAC | 282.79 | WPQ | 196.18 |
| WRP | 741.50 | WAG | 352.16 | WRH | 277.20 | WVE | 192.50 |
| WPC | 739.03 | WVD | 343.00 | WVV | 275.33 | WAH | 189.55 |
| WAR | 664.68 | WVA | 341.25 | WAA | 274.53 | WVF | 188.33 |
| WRW | 603.14 | WVG | 340.63 | WRG | 269.85 | WPW | 182.96 |
| WPS | 588.31 | WVT | 335.63 | WRS | 265.09 | WRE | 160.15 |
| WRT | 555.60 | WVK | 331.38 | WAL | 260.10 | WAF | 153.32 |
| WRV | 541.17 | WRI | 330.4 | WRF | 257.07 | WRL | 150.75 |
| WRA | 534.67 | WAM | 329.85 | WVH | 256.86 | WPG | 134.98 |
| WPV | 511.47 | WVM | 317.00 | WAV | 249.79 | WPN | 128.59 |
| WPD | 480.61 | WPH | 314.57 | WVS | 247.00 | WAY | 117.40 |
| WAE | 461.79 | WRN | 313.20 | WVW | 238.63 | WAW | 103.66 |
| WPL | 461.32 | WPY | 303.79 | WVC | 231.78 | WPI | - |
| WAP | 440.97 | WPP | 303.49 | WPM | 218.75 | WPK | - |
| WPT | 435.08 | WAN | 301.78 | WVY | 214.88 | WAK | - |
| WAD | 432.65 | WRY | 296.50 | WRC | 209.64 | WAS | - |
| WVP | 402.00 | WAT | 296.33 | WPR | 208.94 | WRK | - |
| WVL | 398.88 | WAQ | 289.19 | WPA | 206.01 | WRR | - |
| WVN | 372.88 | WPF | 285.91 | WRD | 203.64 | WVI | - |
| WVQ | 366.00 | WRQ | 285.50 | WAI | 198.88 | WVR | - |

2 IC<sub>50</sub> : half maximal inhibitory concentration.
3 - : not detected within 1

### A proposed strategy to design DPP-IV-IPs based on the repeating dipeptide unit X-proline

Recently, the development of protein synthesis technology has enabled screening of DPP-IV inhibitory activity among chemically synthesized peptides. These studies were conducted based on the property of preferential cleaving of X-proline or X-alanine from the N terminus of peptides by DPP-IV. However, the feasibility of designing the repeating dipeptide unit X-proline remained unclear. To address this problem, we used our model to investigate the potential of this design strategy.

As VPI has the highest hDPP-IV inhibitory activity^27^, the dipeptide unit VP was selected for further study. A library of 20 pentapeptides containing two VP repeats (VPVPX) was designed, and the inhibitory activity of these 20 kinds of pentapeptides were predicted by our model. In addition, IPIPI and IPVPI were also predicted by our model, as IPI has the highest porcine DPP-IV inhibitory activity^27^. The model predicted that 14 out of 22 pentapeptides exhibit inhibitory activity against DPP-IV, which has provided us with new insights into the properties of pentapeptides capable of inhibiting DPP-IV (Supplementary Table 4).

To verify the properties of the peptides predicted, these peptides were chemically synthesized, and their IC_50_ were measured (Fig. 5, Supplementary Table 5). Eight pentapeptides were identified as false negatives, whereas the IC_50_ for the other peptides were correctly predicted. The top three peptides with the highest IC_50_ were IPIPI, VPVPH, and VPVPC, which were 47.47 µM, 51.11 µM, and 54.77 µM, respectively. Inhibitory activity of IPIPI was 2.18-fold higher than WAW, which is the tripeptides with highest inhibitory activity detected in this study. This result indicates that pentapeptides with the repeating dipeptide unit VP exhibit higher DPP-IV inhibitory activity as compared with tripeptides and thus suggests that designing peptides with repeating units is a more efficient strategy.

**Fig. 5.**
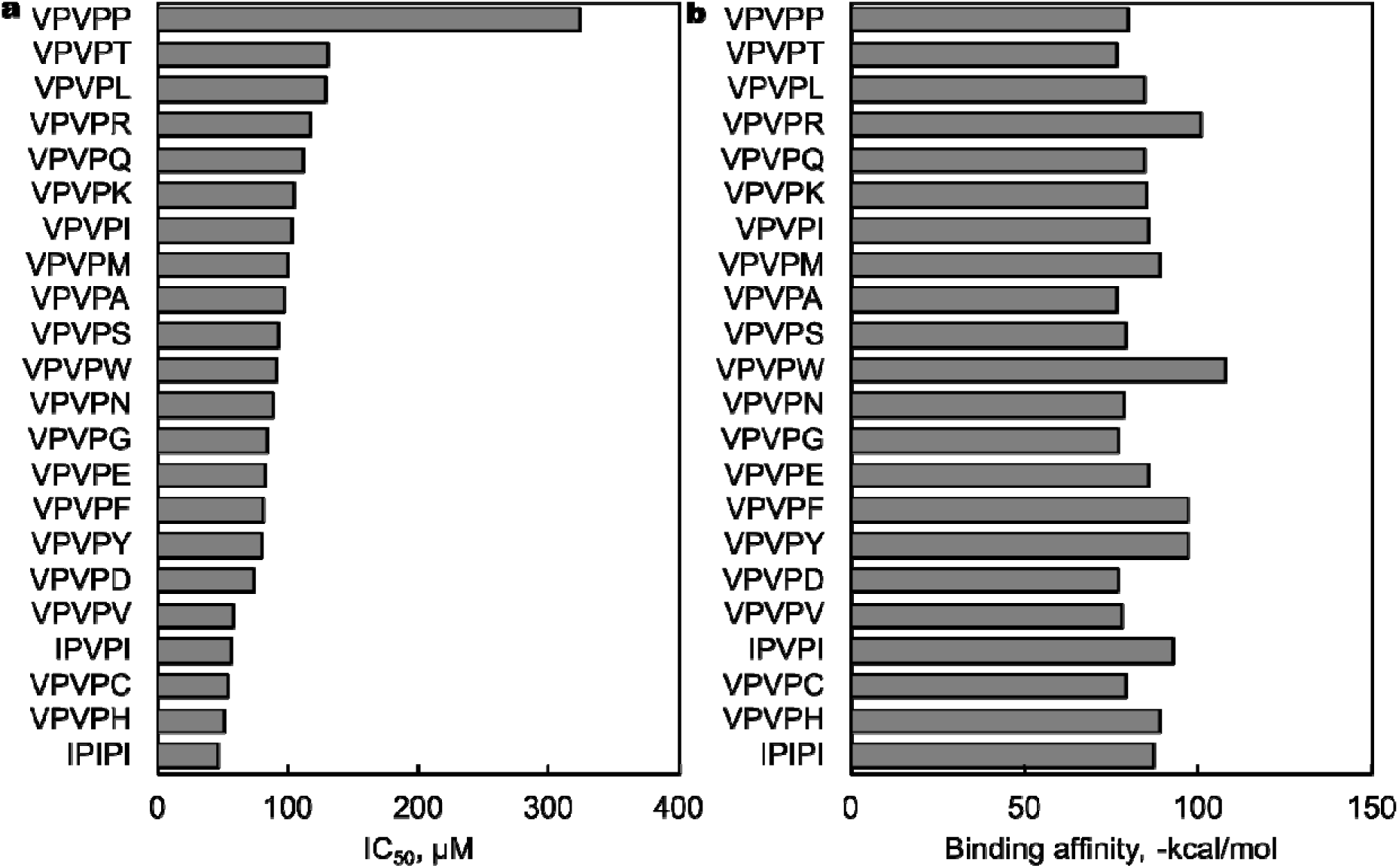
Characteristics of the activity of DPP-IV inhibitory pentapeptides. The pentapeptides were characterized by (a) IC_50_ and (b) binding affinity.

To understand the DPP-IV inhibitory activity of these pentapeptides, their binding energy, which is inversely proportional to inhibitory activity, was analyzed with MDockPeP^28^. We calculated the binding affinities of pentapeptides to hDPP-IV and found that VPVPW, VPVPR, and VPVPY had the lowest affinity. However, a difference was observed in their ranking with respect to binding affinity relative to that for IC_50_ (Supplementary Table 5). This difference might have resulted from (1) the docking method, which reflects only the first step of the interaction between the polypeptide and DPP-IV and cannot reflect the subsequent two consecutive VP cleavage steps, or (2) the docking affinity being calculated based on DPP-IV in the static state, which thus did not consider the dynamic of DPP-IV.

These results suggested that a design strategy based on repeating X-proline units is effective and feasible, although the efficiency of peptides with repeating units of X-alanine should be investigated.

## Discussion

In this study, we have successfully built a PLM-based model BERT-DPPIV for mining novel DPP-IV-IPs and applied it to guide biological experiments for screening and design. Our model, BERT-DPPIV, had demonstrated Acc of 0.894, which is higher than those previously reported, e.g., StackDPPIV^16^. Furthermore, this model can also capture more biologically relevant information from peptide sequences, notably the cleavage site information of DPP-IV and structure information of peptide sequences (Fig. 2, Supplementary Fig. 5). Our model can automatically capture information from various aspects of sequence, structural feature information from the peptide sequence, and the specific cleavage site features from the peptide dataset. This renders our model more advantageous than StackDPPIV, which has been the best model described to date, as the latter must extract sequence and structure feature information by feature engineering.

Although our model has achieved good performance, there is room for further improvement. As BERT models that use the word segmentation method with overlap, e.g., DNABERT^29^, typically show higher accuracy than those that do not, our model which is based on the non-overlapping word segmentation method, can be further improved by using an overlapping segmentation method. The attention-based BERT model adopted in this study overcomes poor interpretability of DL methods. Visualizing the model’s attention during learning revealed what the model learned and how this model had changed during training. Through this strategy, we discovered that our model can capture cleavage site information for DPP-IV in addition to the physicochemical properties and structural information of the peptide sequence (Fig. 2, Supplementary Fig. 5), which had not previously been reported as a classifier. This is the first use of the PLM BERT to discover enzyme cleavage site information from substrate information, and this model should be extremely helpful in understanding the biological implications of DPP-IV. Apparently, this feature can be extended to other enzymes after pre-training with respective databases, and it should be applicable to understanding the functions of different enzymes and to mining the cleavage site information of enzymes.

In contrast with the previous classifiers for inhibiting DPP-IV peptides^14–16^ that were tested only on a dataset and were not verified by actual screening experiments, we used the proposed model to screen tripeptides with DPP-IV inhibitory activity and showed experimentally that our model had an accuracy of 90%. DPP-IV inhibitory activity was identified in 72 out of 80 tripeptides predicted (Table 1). This result suggested that our model can be used to aid in the screening of functional peptides that inhibit DPP-IV.

The number of reported DPP-IV inhibitory polypeptides is still limited due to the absence of strategies for designing DPP-IV inhibitory polypeptide libraries. Therefore, we have also proposed a new peptide design strategy to design polypeptides that inhibit DPP-IV based on repeating dipeptide units to accelerate mining of DPP-IV inhibitory polypeptides. This design strategy has been verified and shown to be feasible through the predictions of our model and subsequent biological assays. Twenty-two pentapeptides containing VP or IP were discovered to inhibit DPP-IV efficiently. This design strategy can help researchers to efficiently explore peptides that inhibit DPP-IV. The accuracy of the model for pentapeptide prediction was 63.6% despite the lack of sequences with repeating dipeptide units in the training dataset. It is likely that the model can be improved by increasing the availability of data through biological assays. Verification of the efficiency of the model in predicting long peptide sequences is in progress.

In conclusion, we have constructed the first PLM-based DPP-IV classifier and have verified its performance experimentally. Furthermore, based on the assistance of the classifier, 72 peptides were revealed to inhibit DPP-IV. A novel design strategy for peptides with DPP-IV inhibitory activity was proposed and verified, based on which 22 pentapeptides were discovered to have DPP-IV inhibitory effects.

In conclusion, we have proposed BERT-DPPIV, a PLM model which is very effective in DPP-IV-IPs mining. This model could aid screening and design of DPP-IV-IPs, which will greatly accelerate the process and reduce the cost of anti-diabetic drug development.

## Methods

### Database

The pre-trained protein data were downloaded from UniProt and can be accessed on Google Drive at https://drive.google.com/file/d/1QeXWV5_OIKgms7u5ShNfvPEKPBLOC3UT/view?usp=sharing. This dataset contains 556,603 protein sequences^30^. The dataset used for fine-tuning was previously described and was used to train and test our proposed model^14^. The benchmark dataset includes 532 DPP-IV-IPs and the same number of peptides with no inhibitory activity against DPP-IV. The number of IPs and non-inhibitory peptides in the independent dataset was 133 peptides each.

### Reagents and peptides

hDPP-IV (>200 U/ml) was obtained from the ATGen company (South Korea). Gly-Pro *p*-nitroanilide (Gly-Pro-pNA) was obtained from the Cayman Chemical Company (Michigan, US). Peptides containing VPVPX, WPX, WAX, WRX, and WVX (where X represents any one of 20 amino acids) with purity > 95% were synthesized by Gen-Script (Suzhou, China).

### Training of the PLM

BERT^31^ is a pre-trained language model developed by Google for natural language text applications and achieves state-of-the-art results on downstream NLP tasks through transfer learning. We pre-trained 12-layer BERT-based language models on 556,603 protein sequences from the UniProt dataset^30^.

To adapt our model to different sequence lengths, we used different word segmentation lengths, kmer = 1, kmer = 2, and kmer = 3, to generate three pre-trained BERT models. The pre-training process consisted of two pre-training tasks: masked language model (MLM) and next sentence prediction (NSP), so that our pre-trained BERT models could capture word-level and sentence-level representations and learn the common features of protein sequences. MLM is able to model complex relationships between amino acids and capture the evolutionary information of proteins^32^. During the MLM task, 15% of the tokens among the original protein sequence were randomly masked to obtain the masked sequence, and a special token [MASK] was used to replace the masked token 80% of the time, a random token was used 10% of the time, and the selected token was unchanged 10% of the time. The model then predicted the masked tokens based on the context of the unmasked sequence for language modeling. For training, we minimized the negative log likelihood of the true amino acid at each of the masked positions using equation (1),

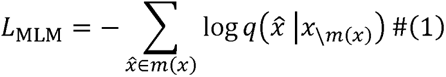

Where *L*_MLM_ is the loss function of MLM, *m*(*x*) and *x_\m(x)_* denote the masked tokens from the protein sequences *x* and the remaining tokens, respectively. During the NSP task, the data were randomly divided into two equal parts A and B: choosing the protein sequence segments A and B for each pretraining example, 50% of the time B is the actual next sentence that follows A (labeled as IsNext), and 50% of the time it is a random sentence from the corpus (labeled as NotNext). BERT was trained by identifying whether these protein sequence segment pairs were continuous. The loss function of the NSP task, *L*_NSP_ was described as equation (2),

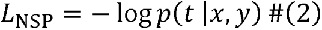

where (*t* = 1 if *x* and *y* are continuous protein segments from protein corpus.

Then MLM and NSP were trained together, with the goal of minimizing the combined loss function of the two strategies as defined in Eq. (3).

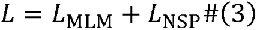

The pre-training hyperparameters included train steps of 10 million times, a learning rate of 2 × 10^−5^ and a batch size of 32 to train the BERT model^30^. After the pre-training process, we obtained three pre-trained BERT models. To construct the PLM for a specific downstream task that identifies and predicts DPP-IV-IPs, we modified the pre-trained BERT models by adding a classification layer on top of the BERT output for the [CLS] token. We fine-tuned the pre-trained model on a benchmark dataset containing DPP-IV-IPs without major architectural modifications. For the fine-tuning hyperparameters, we used a learning rate of 2 × 10^−6^, a batch size of 32, a warm-up proportion of 0.1, and an average training of 50 epochs.

### Performance evaluation of the model

We used four general quantitative indicators to evaluate our model: Acc, sensitivity (Sn), specificity (Sp), and the MCC, each of which is defined by equation (4), equation (5), equation (6), and equation (7), respectively. TP (true positive) is the number of correctly predicted DPP-IV-IPs, FN (false negative) is the number of DPP-IV-IPs that were in fact predicted to be non-DPP-IV-IPs, TN (true negative) is the number of correctly predicted non-DPP-IV-IPs, and FP (false positive) is the number of non-DPP-IV-IPs that were in fact predicted to be DPP-IV-IPs.

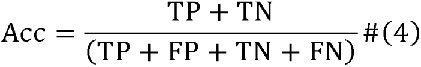

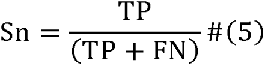

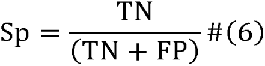

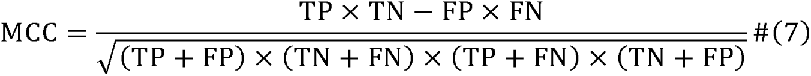

Sn and Sp reflect the model’s ability to recognize DPP-IV-IPs and non-DPP-IV-IPs, respectively, and Acc embodies the overall prediction effect of the model. The value range of the three is [0,1], and the larger the value, the more accurate the model’s prediction. MCC is usually considered a balanced indicator and can be used even if the sample is not balanced. Its value is between –1 and **+**1, reflecting the correlation between the true label of the sample in the testing set and the predicted result. The higher value indicates a greater correlation. When the value is close to **+**1, the classification performance of the model is excellent; when it is close to –1, the prediction result of the model is the opposite of the actual result; and when it is close to 0, the prediction result of the model is similar to a random prediction. In addition, the area under the receiver operating characteristic (AUC) was used as another statistical metric. Considering these five evaluation indicators, the performance of the classification models can be better evaluated.

### Visualization of the Model

Representation visualization was used to analyze the learning capacity of the model. To the original protein sequence, *x* = [*x*_l_,…, *x_n_*] will, we added the special start and end tokens, CLS and SEP, respectively, and got the final input sequence *x*’ = [CLS, *x*_l_,…, *x_n_*, SEP]. The resulting sequence representation *h_i_* was obtained from hidden states in the *i^th^* layer of BERT-DPPIV.

Analyzing the information contained in the representation learned by the model, we used the vector corresponding to CLS in each layer of BERT-DPPIV to represent the analytical sequence and analyzed the representation at the residue level and sequence level by t-SNE. At the residue level, amino acids were divided into six categories, consisting of aromatic (W, F, Y), aliphatic (M, L, I, A, V), positive (R, H, K), negative (D, E), polar neutral (Q, N, S, T), and special-case (G, P, C) residues. At the sequence level, 10 kinds of physicochemical property information were obtained from the modlAMP package^22^.

Attention visualization was used to analyze the information about which the model was specifically concerned and provide interpretable analysis. Each attention head in a model layer produces an attention matrix, α, which indicates the degree of correlation between token pairs. For example, α_*i,j*_ indicates the attention from token *i* to token *j*, and the weight of token *i* for all tokens is 1, as shown in equation (8).

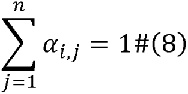

We analyzed attention through a multiscale visualization tool for the Transformer model^24^. The attention-head view in this tool expresses the self-attention matrix a in the form of connected lines and displays the attention patterns generated by one or more attention heads in a given layer. The lines in the head view indicate how much of the hidden state information of the attending token (right) will flow to the attended token (left) (Fig. 2d). The different colors of the lines indicate different attention heads, whereas the color depth of the line is related to the attention weight. The model view in the tool provides a global view of attention patterns across all layers and heads of the model. We then use a slightly simplified version of AlphaFold^26^ to obtain the structure information of the polypeptide sequences and explore the relationship between sites of interaction in polypeptide structures and model attention patterns.

We then used the attention matrix to calculate the importance of each token, *d_j_*, in the peptide sequence as shown in equation (9).

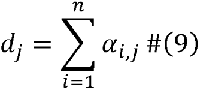

We carried out a statistical analysis of 144 attention patterns including all layers and all heads of each layer. We removed sequences of less than or equal to three amino acids. Then we calculated the importance of all tokens including the special tokens (CLS, SEP) and selected the three most important tokens from each sequence for each attention pattern. We counted the distribution of the top three most important tokens across the different patterns for each sequence with respect to their position and amino acid type. We also selected a specific residue in the sequence and calculated the importance of this residue in 144 attention patterns and visualized the results using a heatmap. We then investigated the interaction between attention and particular amino acids^24^. We used equation (10) for amino acids and defined an indicator function, *f*(*i*,*j*), that returns a value of 1 if this amino acid was present in the token *j* (e.g., if token *j* is a proline).

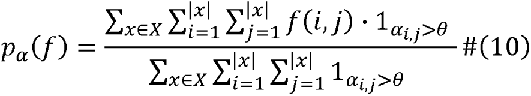

where *p_α_*(*f*) equals the proportion of attention that is directed to the amino acid. We computed the proportion of attention for the fine-tuned model and the pre-trained model directed toward each of the 20 standard amino acids.

### DPP-IV Inhibitory Assay

DPP-IV inhibitory activity was measured according to a slightly modified version of the method described previously^8^. hDPP-IV was used for the triplicate measurements. Peptide solution (20 µl); substrate solution, consisting of 2 mM Gly-Pro-pNA (20 µl); and Tris-HCl buffer (pH 8.0; 20 µl) were pre-mixed in a 96-well microplate. The reaction was initiated by adding 40 µl of hDPP-IV (final concentration, 0.025 U/ml) in 100 µl of the above mixture. The samples were then incubated at 37°C for 60 min. Absorbance at 405 nm was measured using a microplate reader. The negative control mixture included Tris-HCl buffer (pH 8.0), Gly-Pro-pNA, hDPP-IV, and phosphate buffered saline (pH 7.4). The hDPP-IV inhibitory ratio of the sample was calculated according to equation (11),

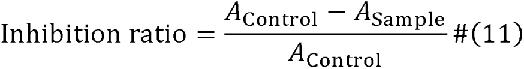

where *A*_Control_ is the absorbance of the control sample with PBS and *A*_Sample_ is the absorbance of the peptide sample.

## Data and Software Availability

The code used in this work is publicly available and can be accessed at https://github.com/guanchangge/BERT-DPPIV. The data used in this work will be made available upon reasonable request and completion of technology transfer agreement.

## Declarations

The authors declare no competing interests

## Author Contributions

X.-H.X., N.Y., Y.L., and C.G. conceived the project and designed the study. C.G. designed the tripeptide and pentapeptide screening experiments; J.C. and C.G. designed the model construction and analysis; C.G. and S.L. performed the tripeptide and pentapeptide screening experiments; C.G. and J.Luo performed model construction; C.G. conducted wet experiments; C.G., J.Luo, and Z.L.T. performed data analysis and interpretation. C.G., J.Luo, J.C., and Z.L.T. drafted and revised the paper. J.Li, Z.W., H.C., Y.W. and C.Z. provided consultation for the project. All authors contributed to the revision of the paper.

## Table of Contents graphic

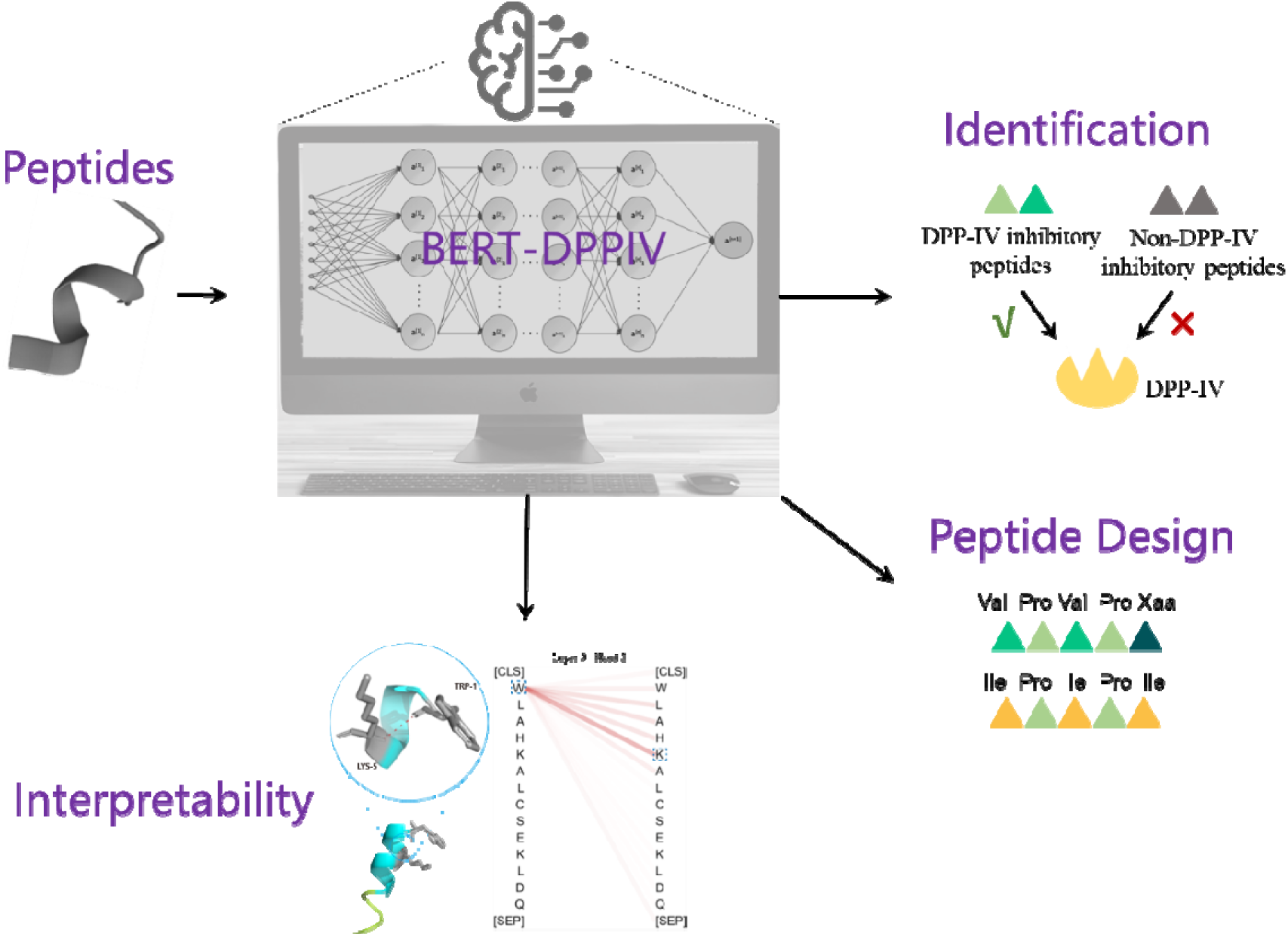

