## Supplement for "Exploration of DPP-IV inhibitory peptide design rules assisted by deep learning pipeline that identifies restriction enzyme cutting site"

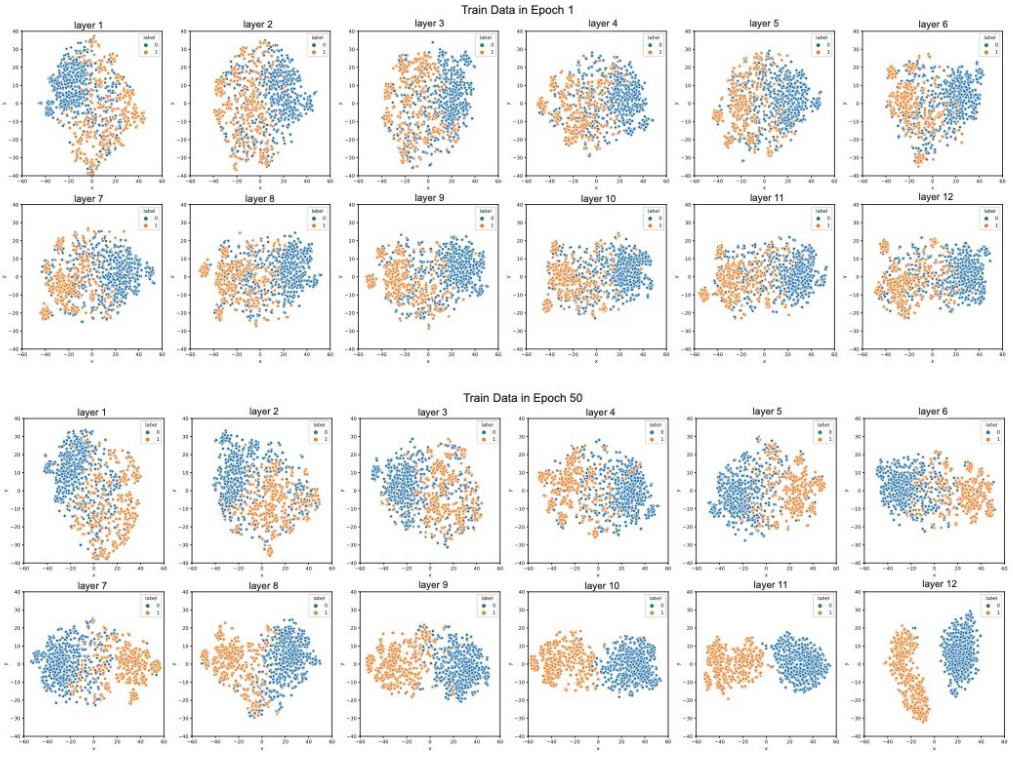


**Supplementary Fig. 1** A visualization of the data representation at the beginning and end of our model training. The vector representation of polypeptides at each layer of the model is dimensionally reduced by t-SNE, and then projected onto a two-dimensional plane. Each cell represents one layer of the model. A detailed visualization of model training can be found in the video.

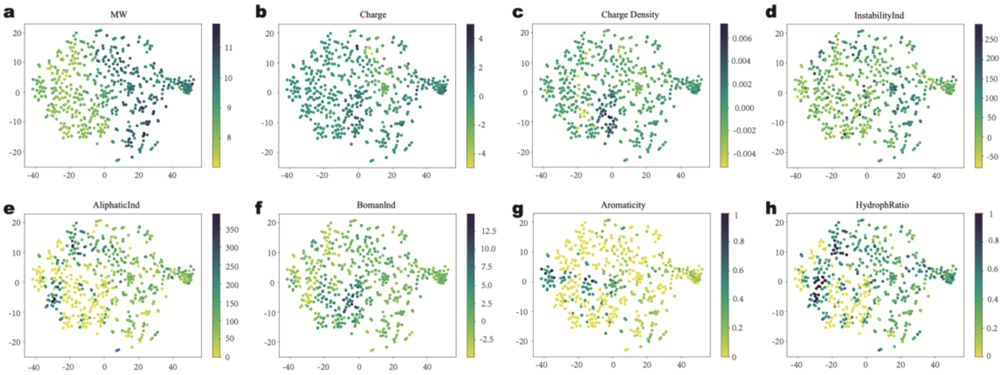


**Supplementary Fig. 2** Analysis of physicochemical properties of sequences based on model-based vector representation. Vector representation of peptide sequence was extracted from the model and then analyzed by t-SNE. The color represents the feature value. (Molecular weight values were logarithmically transformed to base 2)

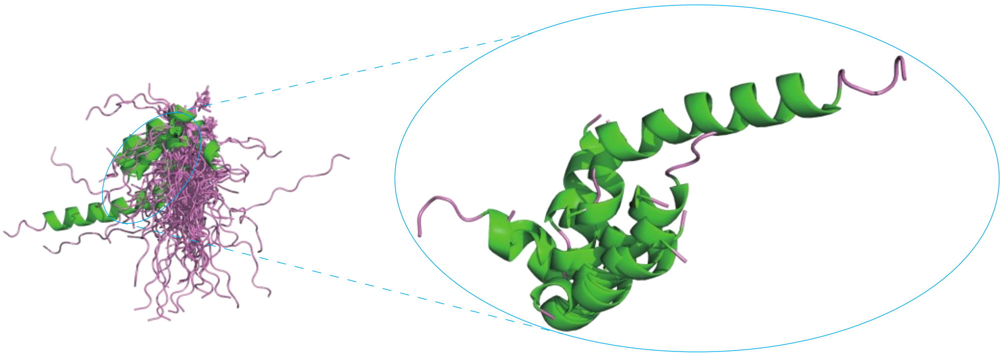

**Supplementary Fig. 3** Structure prediction of peptides. Peptide structure was predicted by APPTEST and displayed by PyMOL. The green and pink represent the alpha helix and loop of the polypeptide sequence, respectively. All structures were displayed (left) and the structure of polypeptides with alpha-helix was displayed (right).

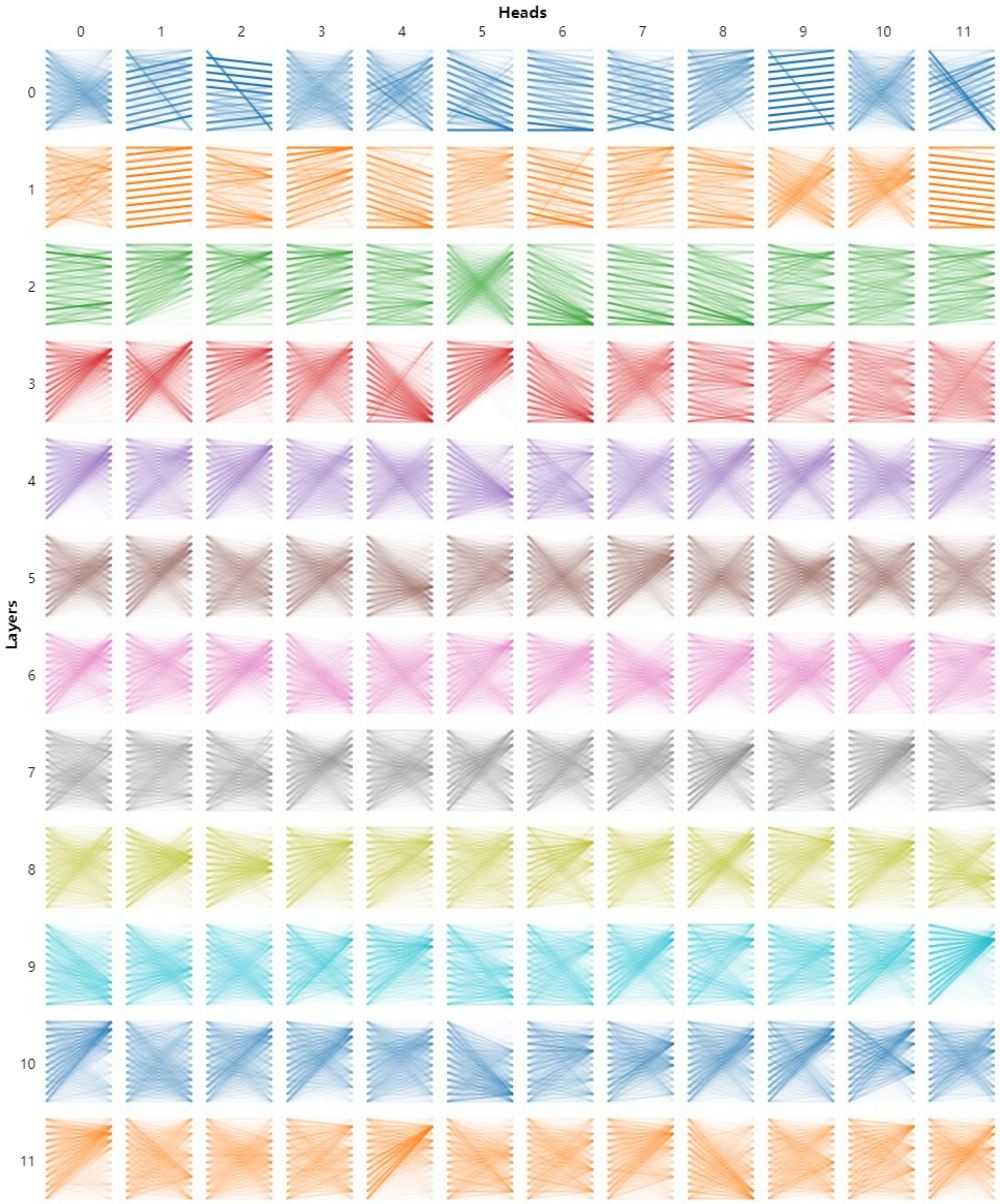


**Supplementary Fig. 4** The attention-head view of YPSKPDNPGE. Each cell in the model view shows the attention pattern for a particular head (indexed by column) in a particular layer (indicated by row). The lines in the head view indicate how much of the hidden state information of the attending token(right) will flow to the attended token(left). The shade of the color represents the size of the attention value.

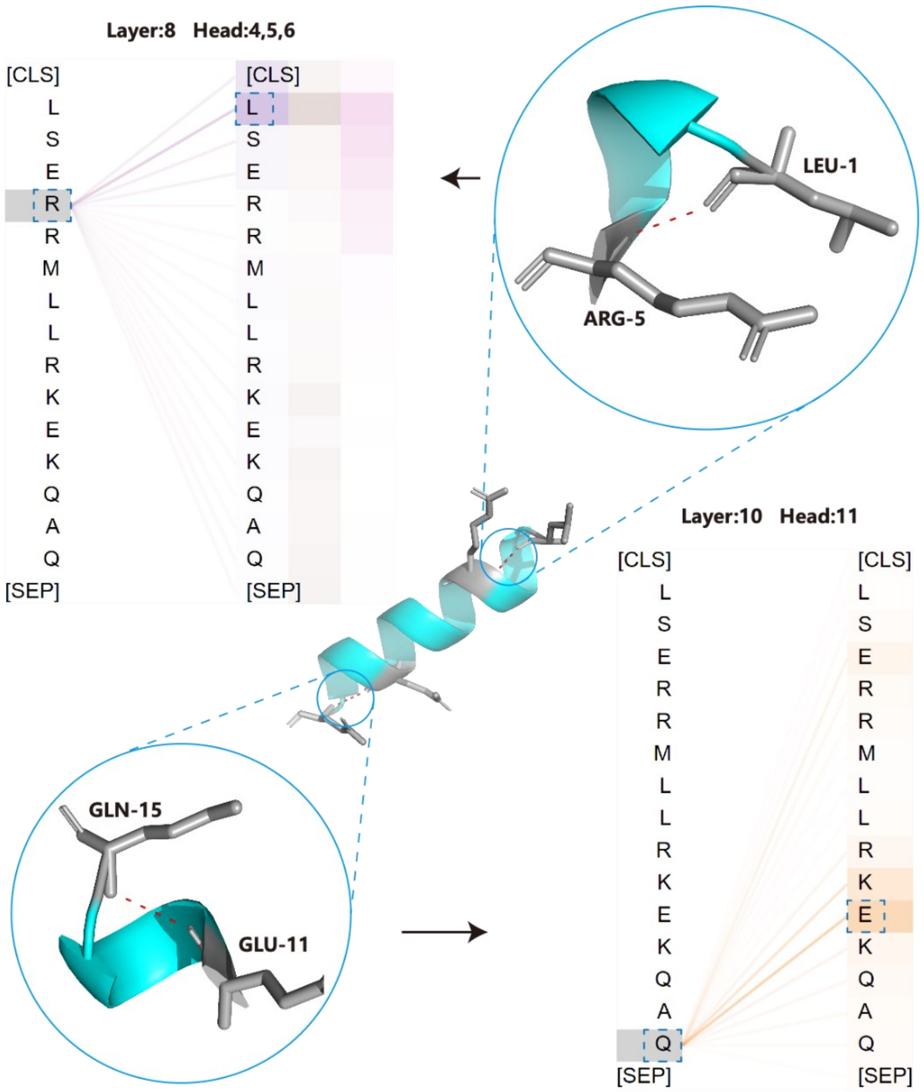


**Supplementary Fig. 5** Attentional visualization of structural information learned by the model. Here, we show how the inner workings of the model's attention heads can be used to analyze a single peptide in more detail. Each attention head performs internally an all-against-all comparison to compute weighted sums for each token over all other tokens in the sequence. High scores indicate that the model learned to put more weight on certain residue pairs (upper and down). The structure of the peptide was shown, GLN-15 can interact with GLU-11 and LEU-1 can interact with ARG-5. When visualizing the attention weights for these sites, we observed that some of the attention heads can capture this interaction information.

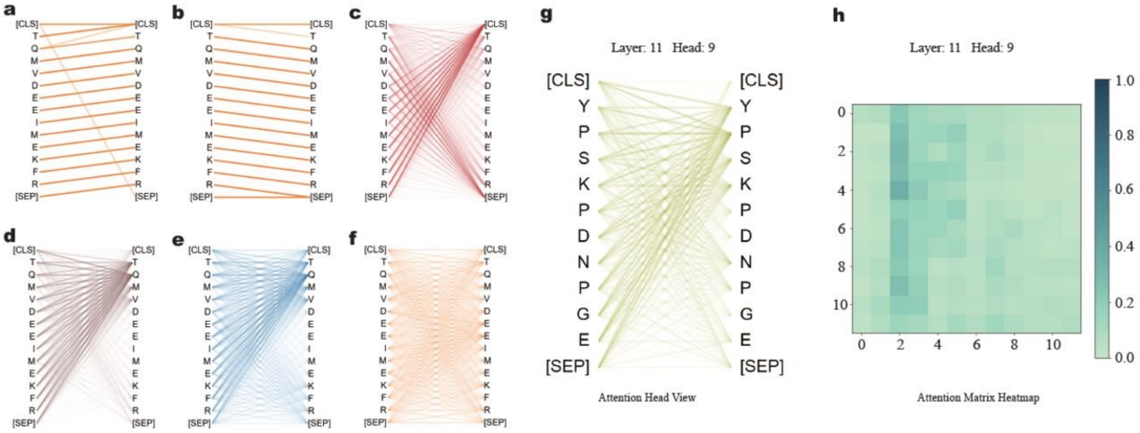


**Supplementary Fig. 6** Visualizing model's attention patterns. Previous-word attention pattern: virtually all the attention is focused on the previous word in the input sequence (a). Next-word attention patterns: virtually all the attention is focused on the next word in the input sequence (b). Delimiter-focused attention patterns: most attention is focused on the [SEP] token or [CLS] token (c). Specific-word attention patterns: virtually all the attention is focused on the specific site (d and e). Attention Related-words attention patterns: attention is paid to identical or related words (f). The attention mainly foucus on the 2th and 3th position of peptide sequence and the corresponding heatmap (g and h).

**Supplementary Table 1** The classification results of BERT-DPPIV and other predictors

| **Model** | **Acc** | **Sn** | **Sp** | **MCC** | **AUC** |
|---|---|---|---|---|---|
| Model with kmer = 1 | 0.891 | 0.857 | 0.925 | 0.784 | 0.957 |
| Model with kmer = 2 | 0.887 | 0.880 | 0.895 | 0.775 | 0.961 |
| Model with kmer = 3 | 0.842 | 0.850 | 0.835 | 0.684 | 0.941 |
| BERT-DPPIV | 0.894 | 0.872 | 0.917 | 0.790 | 0.960 |
| iDPPIV-SCM | 0.797 | 0.789 | 0.805 | 0.594 | 0.847 |
| SVM | 0.865 | 0.874 | 0.856 | 0.731 | 0.939 |
| StackDPPIV | 0.891 | 0.857 | 0.925 | 0.784 | 0.961 |

Acc, accuracy; Sn, sensitivity; Sp, specificity; MCC, Matthews correlation coefficient; AUC, area under receiver operating characteristics curve

**Supplementary Table. 2** Amino acids and the corresponding maximally attentive heads in the pre-trained model and fine-tuned model. The differences between the attention percentages for our model and the background frequencies of each amino acid are significant. The bolded numbers represent the higher of the two values between the pre-train model and the fine-tuned model.

| Code | Name | Background (%) | Pre-trained model | | Fine-tuned model | |
|---|---|---|---|---|---|---|
| | | | Top head | Attention (%) | Top head | Attention (%) |
| A | Alanine | 5.01 | 4-6 | 8.05 | 10-12 | 21.89 |
| R | Arginine | 2.97 | 8-4 | 10 | 6-5 | 20 |
| N | Asparagine | 3.51 | 6-4 | 25 | 10-8 | 11.11 |
| D | Aspartic acid | 3.11 | 11-2 | 10 | 11-11 | 15.71 |
| C | Cysteine | 0.56 | 11-2 | 20 | 9-9 | 14.17 |
| Q | Glutamine | 5.7 | 8-8 | 14.28 | 12-3 | 33.33 |
| E | Glutamic acid | 5.52 | 9-11 | 9.09 | 6-10 | 22.22 |
| G | Glycine | 6.06 | 4-1 | 16 | 6-10 | 14.81 |
| G | Histidine | 2.22 | 11-1 | 8.87 | 12-3 | 66.67 |
| I | Isoleucine | 5.20 | 4-2 | 11.53 | 12-6 | 21.62 |
| L | Leucine | 9.46 | 6-10 | 100 | 6-8 | 26.66 |
| K | Lysine | 3.51 | 11-5 | 13.04 | 12-11 | 14.81 |
| M | Methionine | 3.16 | 9-6 | 7.69 | 12-2 | 15.90 |
| F | Phenylalanine | 4.74 | 10-8 | 15.33 | 8-12 | 20 |
| P | Proline | 17.7 | 5-1 | 50 | 11-9 | 37.93 |
| S | Serine | 3.34 | 8-9 | 28.12 | 10-9 | 26.82 |
| T | Threonine | 4.18 | 11-8 | 16.66 | 12-4 | 22.58 |
| W | Tryptophan | 2.39 | 2-11 | 6.18 | 11-10 | 6.25 |
| Y | Tyrosine | 3.45 | 9-10 | 9.09 | 10-3 | 22.03 |
| V | Valine | 8.01 | 7-8 | 50 | 12-8 | 33 |

**Supplementary Table. 3** The predicted results of tripeptides inhibiting DPP-IV by BERT-DPPIV

| Peptide | Prediction | Peptide | Prediction | Peptide | Prediction | Peptide | Prediction |
|---|---|---|---|---|---|---|---|
| WPA | 1 | WAA | 1 | WRA | 1 | WVA | 1 |
| WPC | 1 | WAC | 1 | WRC | 1 | WVC | 1 |
| WPD | 1 | WAD | 1 | WRD | 1 | WVD | 1 |
| WPE | 1 | WAE | 1 | WRE | 1 | WVE | 1 |
| WPF | 1 | WAF | 1 | WRF | 1 | WVF | 1 |
| WPG | 1 | WAG | 1 | WRG | 1 | WVG | 1 |
| WPH | 1 | WAH | 1 | WRH | 1 | WVH | 1 |
| WPL | 1 | WAL | 1 | WRL | 1 | WVL | 1 |
| WPM | 1 | WAM | 1 | WRM | 1 | WVM | 1 |
| WPN | 1 | WAN | 1 | WRN | 1 | WVN | 1 |
| WPP | 1 | WAP | 1 | WRP | 1 | WVP | 1 |
| WPQ | 1 | WAQ | 1 | WRQ | 1 | WVQ | 1 |
| WPT | 1 | WAT | 1 | WRT | 1 | WVT | 1 |
| WPV | 1 | WAV | 1 | WRV | 1 | WVV | 1 |
| WPW | 1 | WAW | 1 | WRW | 1 | WVW | 1 |
| WPY | 1 | WAY | 1 | WRY | 1 | WVY | 1 |
| WPI | 1 | WAI | 1 | WRI | 1 | WVI | 1 |
| WPK | 1 | WAK | 1 | WRK | 1 | WVK | 1 |
| WPR | 1 | WAR | 1 | WRR | 1 | WVR | 1 |
| WPS | 1 | WAS | 1 | WRS | 1 | WVS | 1 |

1: represents that this peptide can inhibit DPP-IV
0: represents that this peptide cannot inhibit DPP-IV

**Supplementary Table. 4** The predicted results of pentapeptides inhibiting DPP-IV

| Pentapeptide | Prediction | Pentapeptide | Prediction |
|---|---|---|---|
| VPVPA | 1 | VPVPM | 1 |
| VPVPC | 1 | VPVPN | 0 |
| VPVPD | 1 | VPVPP | 0 |
| VPVPE | 1 | VPVPQ | 0 |
| VPVPF | 1 | VPVPR | 1 |
| VPVPG | 0 | VPVPS | 1 |
| VPVPH | 0 | VPVPT | 1 |
| VPVPI | 0 | VPVPV | 1 |
| VPVPK | 1 | VPVPW | 1 |
| VPVPL | 1 | VPVPY | 1 |
| IPIPI | 0 | IPVPI | 0 |

**Supplementary Table. 5** The characteristic results of pentapeptides inhibiting DPP-IV by biology experiment and docking.

| Pentapeptide | $IC_{50}$ (μM) | Affinity (kcal/mol) | Pentapeptide | $IC_{50}$ (μM) | Affinity (kcal/mol) |
|---|---|---|---|---|---|
| VPVPA | 98.05 | -76.4 | VPVPM | 99.92 | -89.2 |
| VPVPC | 54.77 | -79.4 | VPVPN | 88.74 | -78.8 |
| VPVPD | 74.5 | -77.4 | VPVPP | 323.7 | -80.1 |
| VPVPE | 83.44 | -85.7 | VPVPQ | 111.55 | -84.8 |
| VPVPF | 81.87 | -97 | VPVPR | 117.71 | -101.2 |
| VPVPG | 84.91 | -77.3 | VPVPS | 92.87 | -79.3 |
| VPVPH | 51.11 | -88.8 | VPVPT | 130.67 | -76.7 |
| VPVPI | 103.19 | -85.6 | VPVPV | 59.05 | -78.2 |
| VPVPK | 104.29 | -85.1 | VPVPW | 92.1 | -108.2 |
| VPVPL | 129.62 | -84.6 | VPVPY | 80.73 | -97.3 |
| IPIPI | 47.47 | -87.4 | IPVPI | 56.98 | -93 |
