## Supplementary Methods for "Exploration of DPP-IV inhibitory peptide design rules assisted by deep learning pipeline that identifies restriction enzyme cutting site"

### A. Model

In the Natural Language Process (NLP) models, the pre-trained models have many advantages in contrast to directly training the model to complete the target task, they can leverage the large unlabeled datasets to learn universal representations of data, which are beneficial for various downstream tasks and can avoid training a new model from scratch. Therefore, the pre-train strategy was adapted for constructing our model and the deep bidirectional language representation (BERT) model architecture with 12 layers, 768 hidden, and 12 heads was used as the initial model for pre-training that was the first to be proposed by Google AI Language[1]. Then we take the 556,603 protein sequences from the UniProt database integrating the data of SWISS-PROT, TrEMBL, and pir-PSD as pre-train data to pre-train the BERT model[2]. To adapt our model to different sequence lengths, different word segmentation was adapted including kmer=1, kmer=2, and kmer=3 to get three kinds of pre-trained BERT models. The pre-train process contains two pre-training tasks: masked language model (MLM) and next sentence prediction (NSP) so that our pre-trained BERT models can capture word-level and sentence-level representations and learn the common features of protein sequences.
The BERT model is composed by the Bert Encoder and Bert Embedding.

#### A.1 Bert Embedding

To help the model distinguish between the two sentences in training, the input is processed in the following way before entering the model:
A [CLS] token is inserted at the beginning of the first sentence and a [SEP] token is inserted at the end of each sentence.
A sentence embedding indicating Sentence A or Sentence B is added to each token.
A positional embedding is added to each token to indicate its position in the sequence. The concept and implementation of positional embedding are ref. [1]
Therefore, the input embedding consists of three parts, which are token embedding, segment embedding, and position embedding (Supplementary Fig. 7)

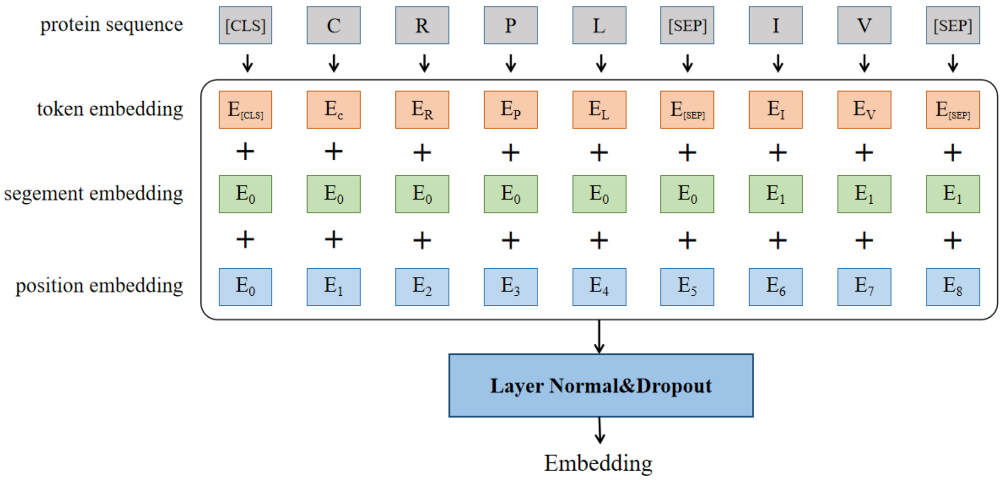


**Supplementary Fig. 7** The architecture of Bert embedding

##### Token embedding

The input is the id number of the token:

$$tokens = [b \times l]$$

Where b is the batch size and l is the sequence length.
The input will be converted to token embedding through the learnable embedding function.

$$X_{token\_embedding} = [b \times l \times d_{model}]$$

##### Segment embedding

Segment embedding is used to distinguish two different sequences. Using the embedding of the learnable embedding function to represent different sentences, the encoded vectors of tokens located in the same sentence are consistent.

$$X_{segment\_embedding} = [b \times l \times d_{model}]$$

##### Positional embedding

Positional embedding is used for self-attention mechanisms that cannot capture the positional information of the sequence. In the original Transformer structure, the sine and cosine functions are used to encode different position information, and the learnable embedding function is used in Bert to represent different positions:

$$X_{position\_embedding} = [b \times l \times d_{model}]$$

After position encoding, it is equivalent to getting three array $X_{token\_embedding}$, $X_{segment\_embedding}$, $and\ X_{position\_embedding}$ that have the same dimensions, when they are added together, get a new word embedding containing position and sentence information.

$$X_{embedding} = X_{token\_embedding} + X_{segment\_embedding} + X_{position\_embedding}$$

The obtained embedding finally passes through the Layer-normalization and dropout functions to get the input of the BertEncoder part

#### A.2 BERT Encoder

The BERT Encoder contains three different modules: Multi-Head Attention, Feedforward, and Layer norm & dropout (Supplementary Fig. 8)

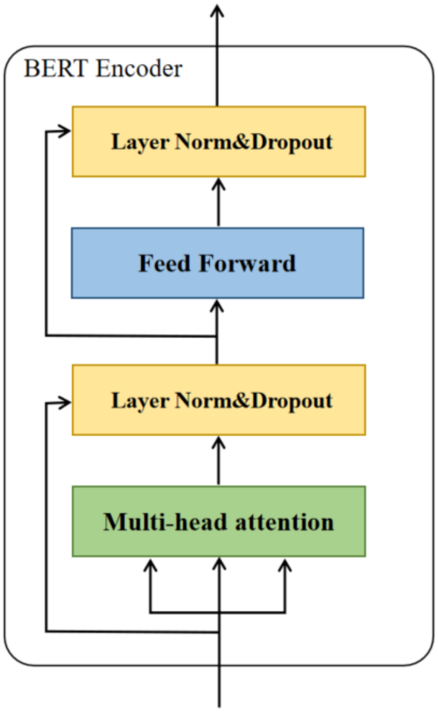


**Supplementary Fig. 8** The architecture of BERT Encoder

##### Multi-Head Attention

Here we first introduce the concepts of Query, Key and Value. Query is the meaning of query, and Key is the key used to compare with the Query you want to query, and then get a score (relevance or similarity) by multiplying the value by Value to get final result. The dimensions of Q, K, V are consistent with $X_{embedding}$. It is generated from the input of oneself $X_{embedding}$, that is, a linear mapping is performed to generate Q, K, V:

$$Q = X_{embedding} * W^Q$$
$$K = X_{embedding} * W^k$$
$$V = X_{embedding} * W^v$$

Where $W^Q$, $W^k$, and $W^v$ were the weight matrices.

After obtaining the matrices Q, K, and V, the output of Self-Attention can be calculated (Supplementary Fig. 9). The calculation formula is as follows:

$$Attention(Q, K, V) = softmax(\frac{QK^T}{\sqrt{d_k}})V$$

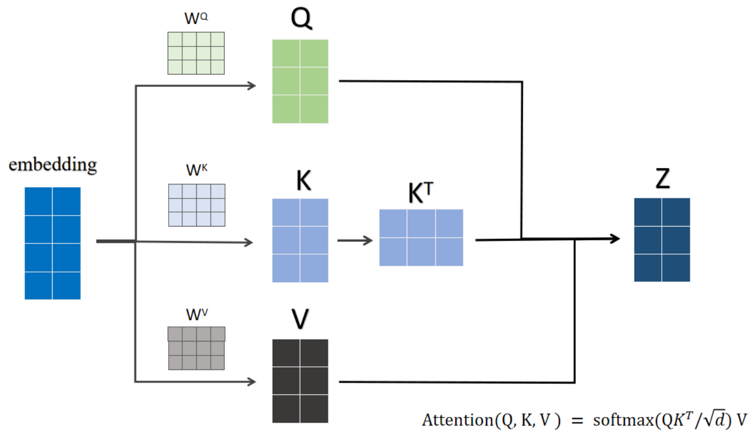


**Supplementary Fig. 9** The calculation of self-attention

where $d_k$ is the number of columns of the Q and K matrix, i.e. the vector dimension. Using a single-head attention mode, the model may not be able to pay attention to a variety of different information, use a multi-head attention mechanism to enhance the model's representation ability. The calculation formula is as follows:

$$MultiHead(Q,K,V) = Concat(head_1, head_2, \cdots, head_h)W^O$$

$$MultiHead(Q,K,V) = softax(\frac{QK^T}{\sqrt{d_k}})V$$

$$head_i = Attention(QW_i^Q, KW_i^K, VW_i^V)$$

where $W_i^Q$, $W_i^K$,and $W_i^V$ were the parameter matrices, $W^O$ is weight matric.

#### Feedforward

The Feedforward layer is a two-layer fully connected layer. The activation function of the first layer is Relu, and the activation function of the second layer is not used. The corresponding formula is as follows:

$$\mathrm{Feedforward(X)} = \max(0, \mathrm{XW}_1 + b_1) + b_2$$

where X represents the input, $\mathrm{W}_1$represent parameter matric, $b_1$ and $b_2$ represent the parameter values.

#### Layer Norm & Dropout

Here are mainly two operations Layer-normalization and Dropout, before normalization, the representation of the current output is added to the previous representation for residual connection. the formula is as follows:

$$y = \mathrm{Dropout(LayerNorm(X + F(X)))}$$

where X represents the input of the F function and the F function is the Multi-head attention or feedforward layer.

The output of the Multi-head attention and feedforward layer will input the Layer Norm &Dropout

$$\mathrm{LayerNorm(X + Multihead\ Attention(X))}$$

$$\mathrm{LayerNorm(X + Feedforward(X))}$$

#### A.3 pre-training process

In the pre-training process, two kinds of tasks were performed including masked language modeling (MLM) and next sentence prediction (NSP). The input amino acid sequence was divided into different tokens using different tokenization methods and add the CLS token before the starting position of the sequence and the SEP token is added between the sequences to obtain the model input vector $X = [x_0, x_1, x_2 \cdots x_T]$, its joint probability

$$q(x_{1:T}) = \prod_{t=1}^{T} q(x_t | x_{0:t-1})$$

where $x_0$ is a special CLS token, $x_i$ represents the i-th token.
The conditional probability $q(x_t \, | x_{0:t-1})$ can be modeled by a probability distribution over the vocabulary given linguistic context $x_{0:t-1}$. The context $x_{0:t-1}$ is modeled by neural encoder function $E_\phi(\cdot)$ parameterized with $\phi$ , and the conditional probability is

$$q(x_t \, | x_{0:t-1}) = LM(E_\phi(x_{0:t-1}))$$

where $LM(\cdot)$ is prediction layer function.
And Loss functions of MLM task is：

$$L_{MLM} = -\sum_{\hat{x} \in m(x)} \log \, q(\hat{x} \, | x_{\backslash m(x)})$$

$m(x)$ and $x_{\backslash m(x)}$ denote the masked words from X and the rest words respectively.

Given the pre-train data set, we can train the entire network with maximum likelihood estimation (MLE). Due to [MASK] token does not appear during fine-tuning, a mismatch between pre-training and fine-tuning is created. To mitigate this, [MASK] is not always used to replace the masked word. The training data generator chooses 15% of the token positions at random for prediction. [MASK] is used to replace the masked word 80% of the time, a random word is used in 10% of the time and the selected word keeps unchanged 10% of the time.

In the NSP task, the data are randomly divided into two parts. In 50% of the data, the sentence pairs are contextually continuous, while the remaining half are not. BERT is trained by identifying whether these sentence pairs are continuous. And Loss functions of the NSP task is

$$L_{NSP} = -\log \, p(t \, | x, y)$$

Where $t \, = \, 1$ if $x$ and $y$ are continuous segments from the corpus.

Then MLM and NSP are trained together, with the goal of minimizing the combined loss function of the two strategies.

$$L_{Total} = L_{NSP} + L_{MLM}$$

We take parameters including train steps of 10 million times, a learning rate of $2e^{-5,}$ and each batch size of 32 to train the BERT model [2]. It uses a two-way Transformer as an encoder and performs joint training through two tasks: MLM and NSP so that the model can capture many sequence facts related to downstream tasks. Sufficient training enables our model to fully learn the long-term dependence of proteins and the representation of protein sequence.
After the pre-train process, three kinds of pre-trained BERT models were got. In order to construct the peptide language model for a specific downstream task that identifies and predicts DPP-IV inhibitory peptides, we modify the pre-trained BERT model by adding a classification layer on top

of the BERT output for the [CLS] token, so that the modified model can complete the target task. The classification layer is a fully connected layer so that we fine-tune the pre-trained model without major architectural modifications. The benchmark dataset containing DPP-IV inhibitory peptides was used to fine-tune the pre-trained BERT models and the performance is compared with the trained and independently tested model proposed in the paper where the dataset is located. When fine-tuning, we use a learning rate of $2e^{-6}$, each batch size of 32, a warm-up proportion of 0.1, and an average training of 50 epochs. All codes, data, and models can be found on GitHub at https://github.com/guanchangge/BERT-DPPIV.

### B Visualization

#### B.1 Sequence visualization in the process of fine tuning

In order to observe the change process of the representation ability of the model during the fine-tuning process, we visualized the representation of the model at different levels during the fine-tuning process. Same as residues, sequences in the training set will add special tokens (CLS and SEP) before input to the model, and the 12-layer representation of [CLS] of each epoch model during fine-tuning is taken out. The representation of the sequence will be visualized using the TSNE algorithm in python's toolkit openTSNE. Finally, the dimensionality-reduced sequence representation is displayed on a 2D plane and labeled as inhibitory peptides or Non-DPP-IV inhibitory.

#### B.2 Residue visualization

We visualize the representation of 20 natural amino acids in the model. Each amino acid needs to add special token([CLS],[SEP]) of BERT before entering the model(eg. “A” will be processed as “[CLS] A [SEP]”). After entering the model, we take the vector of the last layer [CLS] of the model as the representation of amino acids. Then we use t-SNE for dimensionality reduction of the representation, which maps the representation from a high-dimensional space to a two-dimensional planar space. We use python's toolkit openTSNE to use the TSNE algorithm. The amino acids are marked as aromatic amino acids (W, F, Y), aliphatic (M, L, I, A, V), positive (R, H, K), negative (D, E), polar neutral (Q, N, S, T), special case (G, P, C), six categories[3].

#### B.3 Sequence visualization with biochemical and biophysical properties

We explore whether the representation obtained by the BERT-DPPIV model contains biochemical and biophysical properties. In this part, we take all the inhibitory peptides data and use the fine-tuned model to obtain its representation (the CLS vector of the last layer as the representation of the sequence). 10 kinds of physicochemical properties information got from modAMP[2] package as criteria for the information contained in the representation. After dimensionality reduction by the TSNE algorithm, we display the representation in a two-dimensional coordinate system and mark each sample point with different colors according to different property values.

#### B.4 Attention visualization

We analyze the attention through a multiple scale visualization tool bertviz for the Transformer model[4]. This visualization tool contains three modes which are attention-head view, model view, and neuron view. The attention-head view in this tool expresses self-attention matrix $\alpha$ in the form of lines connecting and displays the attention patterns generated by one or more attention headers in a given layer. The lines in the head view indicate how much of the hidden state information of the attending token(right) will flow to the attended token(left) and different colors of the lines are represented different attention heads while the color depth of the line is related to the attention weight. The model view in the tool provides a global view of attention patterns across all layers and heads of the model. 144 attention heads are displayed in the form of a table, with rows representing layers and columns representing heads.

We use the attention-head view to explore the relationship between the attention learned by the model and the structure information of the peptide sequence. Firstly we used the peptide tertiary structure prediction tool APPTEST[5] to make predictions on DPP-IV inhibitory peptides data. In this step, we screened out the polypeptide sequences with α-helix structure. Then we select three of these polypeptide sequences with alpha helices and use a slightly simplified version of AlphaFold[6] with Colab to get the structure information of the polypeptide sequence. Use pymol to visualize protein tertiary structures and find their interacting residues. We can find that there are some attention patterns that can find information about the interaction sites in the protein structure.

We also use the model view to summarize the attention patterns in the sequence, and we can find that the attention patterns of the sequence include Next-word attention patterns, Previous-word attention patterns, Delimiter-focused attention patterns, Specific-word attention patterns and Attention Related-words attention patterns.

#### B.5 Statistical Analysis of attention matrix

The attention matrix α describes the relationship between token pairs.$\alpha_{i,j}$ indicate how much token j information needs to be used when computing the representation of token i.

$$\sum_{j=1}^{n} \alpha_{i,j} = 1$$

We use the attention matrix to calculate the importance of each token $d_j$ in the peptide sequence. A higher $d_j$ weight value indicates that token j is receiving more attention in the peptide sequence.

$$d_j = \sum_{i=1}^{n} \alpha_{i,j}$$

We make statistics on 144 attention patterns of all layers and all heads of each layer. Firstly, we remove sequences of length less than or equal to three. The remaining sequences are input into the model after adding special tokens ([CLS], [SEP]), and the attention matrix corresponding to each sequence is obtained. Then we calculate the importance of all tokens including special tokens (CLS, SEP), and take out the three most important tokens for each sequence in each attention pattern. We count the first three tokens that appear in each of the 144 attention patterns in a sequence, and then define the token with the total number of the first three as the most important token in the sequence. We counted the distribution of the top three most important tokens in different patterns for each sequence by position and amino acid, respectively. We also selected a specific residue in the sequence and calculated the importance of this residue $d_{residue}$ in 144 attention modes, and visualized it using a heatmap.

##### B.6 Interaction between attention and particular amino acid

We then investigated the interaction between attention and particular amino acids[7]. We use the next formula for particular amino acids, and define an indicator function f(i,j) that returns 1 if this amino acid is present in token j(e.g if the token j is Pro). We set the attention threshold θ to 0.4 to select high-confidence attention.

$$p_\alpha(f) = \frac{\sum_{x\in X}\sum_{i=1}^{|x|}\sum_{j=1}^{|x|} f(i,j)\cdot 1_{\alpha_{i,j}>\theta}}{\sum_{x\in X}\sum_{i=1}^{|x|}\sum_{j=1}^{|x|} 1_{\alpha_{i,j}>\theta}}$$

$p_\alpha(f)$ equals the proportion of attention that is directed to the particular amino acid. We computed the proportion of fine-tuned model's attention to each of the 20 standard amino acids and visualize it by heatmap with rows representing layers and columns representing heads. The shade of color indicates the percentage of attention that satisfies the condition in this attention head for all data. The bar graph on the right represents the maximum value of the attention ratio in each layer.
